# CODARFE: Unlocking the prediction of continuous environmental variables based on microbiome

**DOI:** 10.1101/2024.07.18.604052

**Authors:** Murilo Caminotto Barbosa, João Fernando Marques da Silva, Leonardo Cardoso Alves, Robert D Finn, Alexandre Rossi Paschoal

## Abstract

Despite the surge in data acquisition, there is a limited availability of tools capable of effectively analyzing microbiome data that identify correlations between taxonomic compositions and continuous environmental factors. Furthermore, existing tools also do not predict the environmental factors in new samples, underscoring the pressing need for innovative solutions to enhance our understanding of microbiome dynamics and fulfill the prediction gap. Here, we introduce CODARFE, a novel tool for sparse compositional microbiome-predictors selection and prediction of continuous environmental factors. We tested CODARFE against four state-of-the-art tools in two experiments. First, CODARFE outperformed predictor selection in 21 out of 24 databases in terms of correlation. Second, among all the tools, CODARFE achieved the highest number of previously identified bacteria linked to environmental factors for human data—that is, at least 7% more. We also tested CODARFE in a cross-study, using the same biome but under different external effects (e.g., ginseng field and cattle for arable soil, and HIV and crohn’s disease for human gut), using a model trained on one dataset to predict environmental factors on another dataset, achieving 11% of mean absolute percentage error. Finally, CODARFE is available in five formats, including a Windows version with a graphical interface, to installable source code for Linux servers and an embedded Jupyter notebook available at MGnify - https://github.com/alerpaschoal/CODARFE.

## Introduction

Microbiomes encompass a vast array of microorganisms, including bacteria, viruses, fungi, and protozoa [Gilbert et al., 2018], inhabiting diverse environments including the human body, plants, soils, and even the International Space Station [Novikova et al., 2006]. This microbial diversity has become a focal point of research to understand the role of microbes in health and disease [Johnson and Versalovic, 2012]. Studies have shown that variations in microbial communities can influence the pathophysiology of various diseases [Lavelle and Hill, 2019, Wen et al., 2017]. Consequently, this has led to significant interest in identifying specific microbial biomarkers that can serve as indicators for these conditions, providing valuable insights into both the diseases themselves and the environmental factors associated with them [Damhorst et al., 2021, Suman et al., 2022]. However, comparison between microbiome data coming from two or more samples can present specific challenges owing to the composition variability, hence the need to develop appropriate statistical approaches [Legendre and Gallagher, 2001, Lin and Peddada, 2020].

Analyzing microbial communities typically means measuring the proportions of different microbial taxa present in a sample, however these measurements do not provide absolute abundance values. Instead, the information is conveyed as ratios or percentages relative to the total microbial community in that sample. This compositional structure creates dependencies among the species proportions. If one species’ abundance increases, the proportions of the others must change even if the real abundances of those species remain constant. This is known as the “closure property” of compositional data. Consequently, mathematical correlations between species in microbiome data are not often reflective of direct biological interactions, but rather a result of the mathematical constraints imposed by the compositional measurements [Gloor et al., 2017].

Additionally, data sparsity arises from the presence of rare or low-abundance species that are difficult to detect and quantify [Kurtz et al., 2015], with the risk of false statistical significance if statistical methods are inappropriately applied [Legendre and Gallagher, 2001]. These unique characteristics demand special statistical approaches that consider the specific structure of sparse compositional data instead of applying traditional methods assuming normality and variable independence [Lutz et al., 2022]. Given these complexities, compositional data analysis (CoDA) techniques have emerged as a prominent research focus in the field of microbiome research. One of these techniques is the centered log-ratio (CLR) [Aitchison, 1982] transformation that allows the representation of compositional data in a Euclidean space, making them suitable for widely used statistical analysis. The statistical analysis then can be used to associate the microbiome predictors – in this research being exclusively taxa – to any target variable. In the context of this research, the “target variable” can be any <u>continuous environmental or clinical</u> variable related to the microbiome sample, such as pH and humidity for soil samples, disease index for human samples or temperature and salinity for marine water samples. Furthermore, the use of machine learning (ML) methods has proven effective for associating subsets of microbial communities with an environmental factor, given the capacity of ML methods to handle high-dimensional data and identify complex patterns [Ghannam and Techtmann, 2021]. However, it is crucial to emphasize that the direct application of ML techniques to compositional data without proper transformations can lead to high error rates [Huang et al., 2023].

While there are a growing collection of ML based tools for microbiome data analysis [Chandrasekhar, 2020, Verster et al., 2022, Calle et al., 2023, Susin et al., 2020, Zhang et al., 2022], aiming to identify biomarkers and gain insights into pathologies, a fundamental gap remains: the ability to accurately predict the variable of interest (such as: soil pH or crohn’s disease active index.) in new samples [Wilhelm et al., 2022]. To address this challenge, we present CODARFE (<u>CO</u>mpositional <u>D</u>ata <u>A</u>nalysis with <u>R</u>ecursive <u>F</u>eature <u>E</u>limination), a new tool that combines CoDA methods with a Recursive Feature Elimination (RFE) process and ML. This approach enables the association of microbiome compositional data with a continuous environmental variable. It also integrates a module for missing predictor imputation and enables the prediction of target environmental variables in new samples.

## Material and Methods

### Evaluation datasets

In this study we collected a total of 30 datasets from three data-sources, and for each source a set of analyses were conducted (Figure S1). The called “Group A: Literature” was collected with the aim of evaluating the coefficient of correlation and processing time required by different methods, and for a hold-out prediction evaluation. In total there are 19 datasets, one for each environmental variable [Wilhelm et al., 2022].The “Group B: ML Repo” was collected to serve as a positive control. This is due to preexisting evidence of a relationship between one or more taxa and the sample metadata variable of interest in the original publication associated with the dataset. The validation is achieved by calculating the number of taxa selected by the tools that are also supported by articles as linked to the target variable. It is composed of five human gut datasets obtained from the collection available in the “ML Repo” [Vangay et al., 2019, Yatsunenko et al., 2012, Gevers et al., 2014, Ravel et al., 2011], where two of these datasets pertain to pediatrics’s Crohn disease, with samples collected from the rectum (n=51) and ileum (n=67); one from a study on the infant microbiome (age 0-2 years) (n=49); and two from the study on the vaginal health, which captured pH (n=388) and Nugent scores (n=388). The “Group C: cross studies” was collected with the purpose to evaluate CODARFÈs generalization power and potential failings when training a model on samples from one project and predicting the variable of interest on a different project (from the same biome). Group C consists of two sets of datasets extracted from MGnify [Richardson et al., 2023]. The first set comprises two projects with pH measurements in arable soil under the effect of ginseng field and cattle (Table 2) [Chrǒnáková et al., 2015, Nguyen et al., 2016] (MGnify identifiers: MGYS00000916 and MGYS00001160). The second set involves age measurements in humans from four projects with samples collected from different parts of the digestive system and subjected to various external effects (such as: antibiotic treatment, HIV, etc) (Table 3) [Di Paola et al., 2016, Raymond et al., 2016, Wang et al., 2020, Noguera Julian et al., 2016] (MGnify identifiers: MGYS00000580, MGYS00001175, MGYS00001188, MGYS00001255).

### Data Preprocessing

The data preprocessing consists of two steps. In the first step, predictors whose variance across samples approximate to zero are removed. A value is considered close to zero if it is less than or equal to 1/8 of the mean variance found in the dataset, by this manner, removing columns that are considered poor in information [Ng et al., 2011].

The second step concerns only the target variable, where values are squared rooted and rearranged between 0-100 if its coefficient of variation (CV) is greater than or equal to 0.2, indicating a wide variation, which may suggest the presence of noise [Smith, 1976]. The square root serves to mitigate this data dispersion if required, which has been proven effective for reducing errors in machine learning models [Zheng and Casari, 2018]. Additionally, if the CV is less than 0.2 and the target contains negative numbers, a simple shift is applied to allow the model to predict the target (due to the poisson distribution used in the model, described ahead).

### Selection of CODA transformation and regression method

Subsequently, four machine learning algorithms were evaluated, using two distinct compositional data transformations, namely Hellinger and Center Log Ratio (CLR) [Legendre and Gallagher, 2001]. These transformations were chosen due to their ability to mitigate the inherent issues of compositional data, where the Hellinger transformation is effective for preserving the Euclidean structure of the data [Legendre and Gallagher, 2001], and the CLR transformation for its ability to transform compositional data into a real-valued space where traditional statistical methods can be applied without the risk of generating misleading results [Aitchison, 1986].

The algorithms evaluated were Linear Support Vector Regression [Chang and Lin, 2011], Linear Stochastic Gradient Descent [Chang and Lin, 2011], Huber regression [Owen, 2007], and Theil Sen estimator [Dang et al., 2008], collectively chosen for their ability to return weights corresponding to the importance of each predictor. Each algorithm was tested with a wide range of hyperparameters and applied to each transformation, termed model-hyperparameter-transformation (MHT), resulting in 2270 combinations. Each MHT was evaluated using RFE [Guyon et al., 2002], which removes 1% of the total predictors (columns) with the smallest weights at each iteration, generating up to a hundred trained models.

Four criteria were used to evaluate each MHT: the *R*^2^ adjusted was chosen to determine the correlation strength with the target variable while selecting a concise number of predictors due to its penalization effect, collaborating to reduce the type 1 error during the fit process; BIC was chosen to differentiate models with similar *R*2 and with different predictor quantities, and it also helps keep low type 1 error due its penalization related to the number of predictors; Rooted Mean Squared Error (RMSE) was used in cross-validation to reduce overfitting, being this the most important metric since overfitting has the worst impact in a prediction model [Shalev-Shwartz and Ben-David,2014]; and lastly, the p-value of the F-test is used to ensure the statistical significance of the selected predictors. All metrics were normalized using the “minmax” method (equation 1) and summed to create the model’s score. The selected MHT is the one with the highest model score. Figure 1A illustrates the complete process.

**Figure 1.**
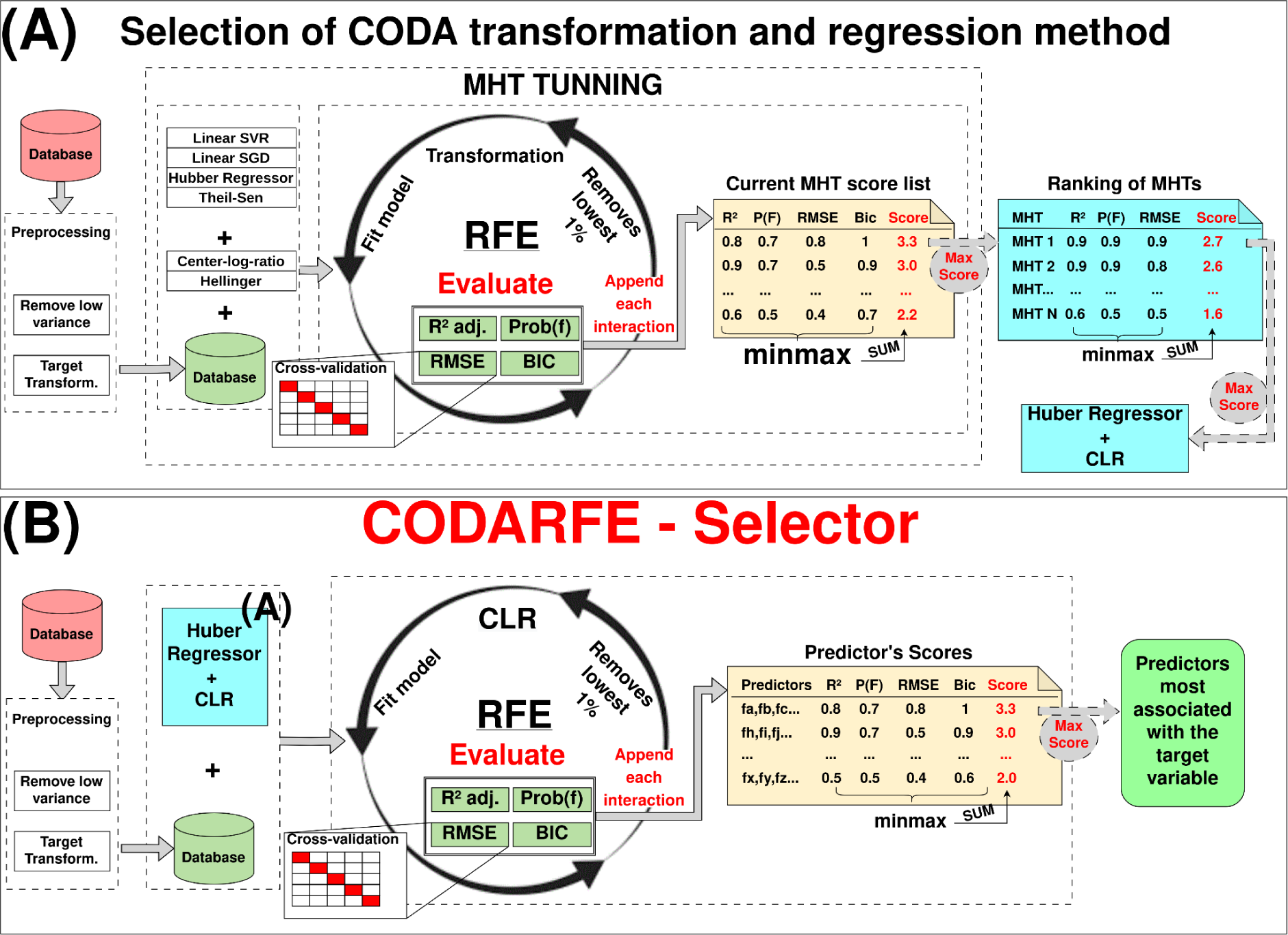
Pipelines from CODARFE-selector and selection of CLR and Huber Regressor as CoDA transformation and regression method.(A) Firstly, CODARFE-selector selects the closely related variables to the target. The data is preprocessed to remove low variation variables, followed by the RFE, in which the trained model is evaluated using four metrics. Finally, the model with the highest “predictors score” (the sum of the “minmax” of the four metrics) is chosen, and the variables used to train the model are stated as the most associated with the target. (B) Subsequently, the selection of CoDA transformation and regression method is performed. The data is preprocessed to remove low variance variables, then RFE is used to train the MHT and evaluate it using four metrics. Finally, the MHT with the highest score (the sum of the “minmax” of the four metrics) is selected.

In the end of the process, the Huber regression yielded the highest scores when paired with the Centered-Log-Ratio (CLR) transformation. The “epsilon” parameter, determining outlier influence, was set to 2, and the “alpha” value, controlling regularization, was set at 0.0003. Unfortunately, the CLR creates an influence of the non-selected predictors into the selected one due to the geometric mean used in its formula. This problem is mitigated with the use of the RFE, where on each interaction, the original data is rescued and re-transformed using only the remaining predictors. Furthermore, the RFE helps with the issues of data dimensionality, which iteratively reduces the number of columns by filtering for importance.

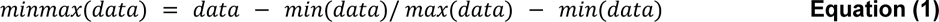

### CODARFE - Selector

The predictor selector is the core of the CODARFE. The results of the selector are the predictors used for correlation to the target and the predictors for the prediction model. It starts with the preprocessing step, then the CLR transformation is applied to the data. The data goes under a Recursive-Feature-Elimination with the Huber regression as an algorithm chosen to give weights (coefficients) to each predictor. The predictors are evaluated following the same process described in the previous section (“Selection of CODA transformation and regression method”), to produce the “Predictors scores”. The set of predictors with the highest score is defined as the predictors most associated with the target variable. Figure 1B depicts the complete process.

### Selection of predictor algorithm

For the predictor algorithm, a new set of regressors were test, namely: Linear Support Vector Regression [Chang and Lin, 2011], Linear Stochastic Gradient Descent [Chang and Lin, 2011], Huber regression [Owen, 2007], Theil-Sen estimator [Dang et al., 2008], Random-Forest [Breiman, 2001], and Support Vector Machine [Chang and Lin, 2011] with various kernels, resulting in 1243 combinations (Figure 2A). Only the CLR transformation was used. Each model was trained on 80% of the dataset and evaluated on 20% with 10 repetitions to calculate the average of the Mean Absolute Error (MAE) estimations (Figure 2A). The Random Forest with parameters “n_estimators = 160” and “criterion = poisson” resulted in a better performance for predictions and was chosen for use in COADRFE.

**Figure 2.**
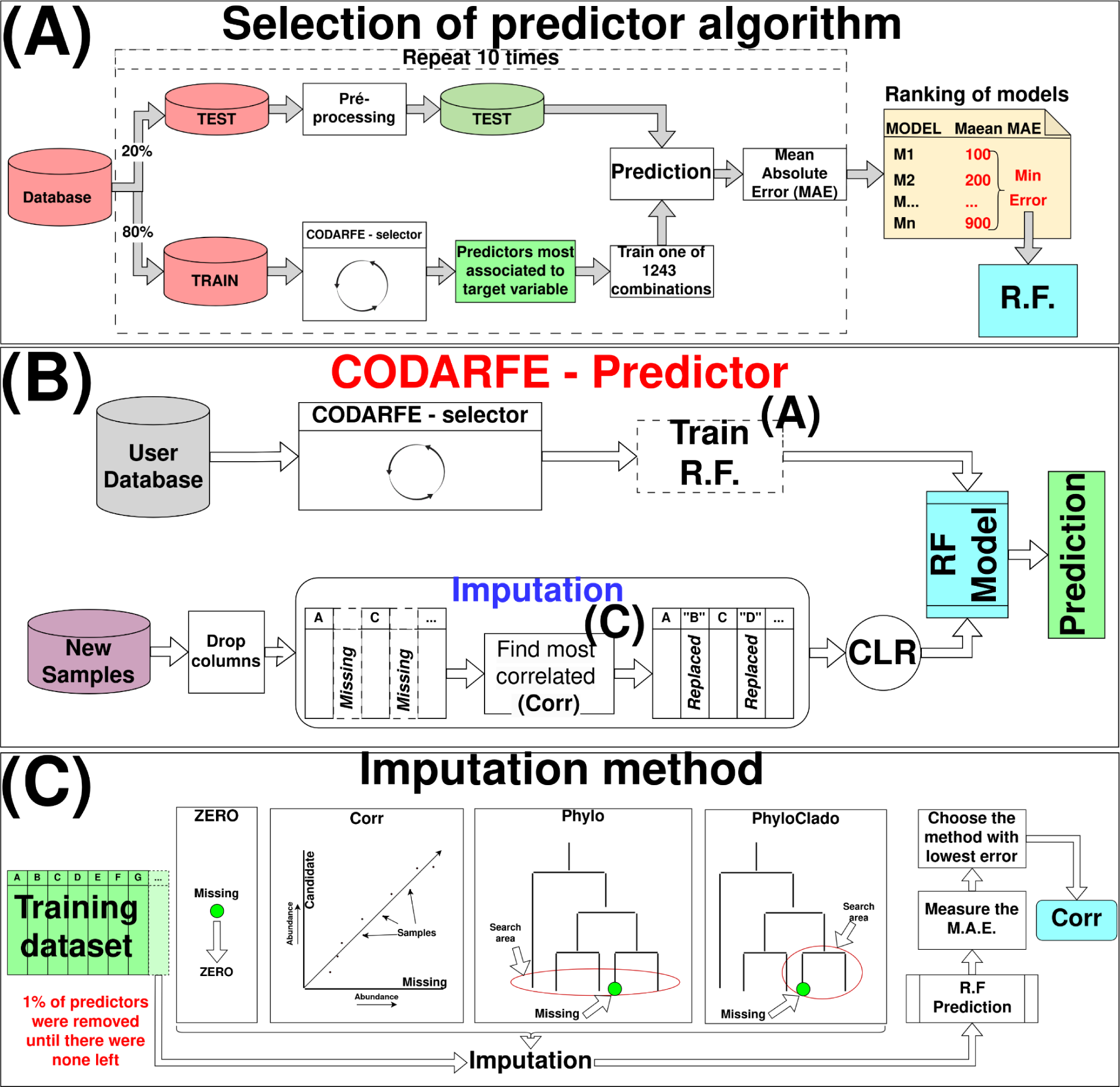
Workflow for CODARFE-Prediction module and the workflows for selecting Random-Forest (RF) as prediction algorithm and correlation (Corr) as the imputation method. (A) A view of the process used to select the Random Forest as a prediction algorithm. First a training database was split so that 80% of the data was used for training and 20% for testing; Then the attributes most associate to target variable were select using the CODARFE-selector; A total of 1243 combinations of ML algorithms and hyper-parameter were tested and the Mean Absolute Error (MAE) was measured with 10 repetitions; As a result, a table of MAEs was created and the algorithm with lowest average MAE was selected as prediction model (RF). (B) The user database is used to select the predictors associated with the target through the CODARFE-selector and then training a Random Forest model. New samples are preprocessed and followed through the imputation step to be used as input for the RF model. The result is the prediction of the new sample’s target variable. (C) The techniques evaluated for replacing (inputting) missing taxa. Four methods were tested in a recursive process where 1% of the total variables from a test dataset was removed on each step simulating increasing missing predictors. Each imputation method was used to replace the missing taxa creating a new dataset for the prediction. This new dataset was used to predict the target and the MAE error was measured. The method with the lowest average MAE was selected as the imputation method.

### CODARFE - Predictor Model

The CODARFE predictor model can only be used after the selector. The selected predictors are used for training a Random Forest algorithm. When new samples are provided for prediction, it first needs to pass through four steps. First predictors (columns) that were not selected are removed from data. Then the data goes through an imputation method that will replace missing predictors by its most correlated available predictor. The Pearson’s correlation on the training CLR-transformed data (CODARFE-selector) is then measured. If a strongly correlated predictor (ρ >= 0.7) is present in the new samples, it will replace the missing one, otherwise the value is set as zero. Finally, the data is CLR-transformed and predicted by the trained Random Forest and the results are obtained (Figure 2B).

### Imputation method

A missing data imputation method was developed to mitigate the issue of dealing with predictors selected by the model which are not present in a new dataset. Four methods were tested. The first method (“zero”) considered the abundance of the missing predictor as 0. The second method (“corr”) created a list containing “K” possible imputation predictors, where each one had a strong correlation (ρ >= 0.7) with the missing one. The Pearson correlation coefficient was calculated from the CLR transformed data, and the value of “K” arbitrarily set to 50. The presence of “K” was to increase the chance that at least one predictor was available in the new samples. The value of the predictor with the highest correlation found in the new sample is used to substitute the value of the missing predictor. However, this method did not ensure the existence of a correlated attribute, returning zero in such cases. The third method (“phylo”) selected the most similar taxa in a phylogenetic tree generated from the union of the new sample with the training set. The distance between two nodes was calculated using the “get-distance” function from the *et3* library in Python version 3.10.6. The selected predictor’s value is assigned in place of the missing predictor. The fourth method (“phyloclade”) was similar to the third method, however it was limited to the clade in which the missing attribute was placed. To test each method, the database ‘aggregate-stability’ present in “Group A: Literature” was used. It was divided in an 80-20 scheme where 80% of the data were used for training a Random Forest and 20% were used for testing. The test data was filtered to contain only the selected predictors. Then it had 1% of its columns removed to simulate missing predictors. After, each method was used to replace the missing ones, and the Random Forest model predicted the results. This process was repeated until there were no predictors left. The Mean Absolute Error was used to choose the imputation method which led to the lowest error rate. Figure 2C illustrates the complete process.

### Comparison against four state-of-art tools

A comparison was done between CODARFE and four other tools in terms of performance on calculating/extracting/evaluating associations between the microbiome and environmental variables. The following describes the tools and how they were used. These tools represent significant advancements in microbiome analysis, each with its advantages and limitations (Table 1). Every tool, including CODARFE, was trained with its default parameters. No fine tuning was performed for any tool to ensure a fair comparison.

**Table 1.** Comparisons of the tools used in this study for performance benchmark. Each tool has its own mode of operation, which is detailed in this table. *Isometric Log-Ratio (ILR). ** Theoretical limit does not exist, but it has a quadratic computational time in relation to the number of columns.

|  | CODARFE | Coda4microbiome | BRACoD | Selbal | CLR-LASSO |
| --- | --- | --- | --- | --- | --- |
| **Filtering method** | RFE | Elastic-Net | MCMC | Balances | LASSO |
| **Transformation** | CLR | ILR* | CLR | ILR* | CLR |
| **Limit of columns** | No | No** | 300 | No** | No |
| **Create graphics** | Yes | Yes | No | No | No |
| **Prediction feature** | Yes | No | No | No | No |

BRACoD employs Bayesian regression models [Verster et al., 2022] to find biological markers of diseases. As recommended by the authors, we used BRACoD with its default settings and filtered the input data to keep only the top 300 most prevalent bacteria. Coda4Microbiome takes into account the inherent variability of count data and uses hierarchical models based on the Bayesian posterior approach [Calle et al., 2023], while Selbal uses the balances method to identify co-abundance patterns among microbiome species [Rivera-Pinto et al., 2018]. For both tools all parameters were set to default, and the “nearZeroSum” function (R package “caret”) was used to filter the input data [Kuhn, 2008]. Finally, CLR-LASSO uses the CLR transformation for Euclidean representation of data and penalized LASSO regression for variable selection [Susin et al., 2020]. Nonetheless it requires the user to set up a lambda value that was defined as the value that mimics the most the number of variables selected by CODARFE.

### Simulating CODARFE’s prediction in identical studies

For the simulations, the datasets from group A were used. For each dataset, 20% of the samples were randomly removed to calculate the mean absolute error (MAE), and 80% were used for model training. This procedure was repeated 10 times, and the average MAE was calculated. The accuracy was measured through the Mean Absolute Percentage Error (MAPE) (equation 2).

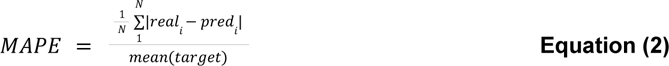

where, “N” is total number of samples, *real_i_* represents the true value in the position “i” and *pred_i_* represents the predicted value in the position “i”.

### Cross-studies prediction simulation

To fairly capture the predictive power, we used the cross-studies approach. We gathered datasets from several projects with different topics, but with at least one environmental variable that could serve as a prediction target included in their metadata. Then we trained the CODARFE - predictor in one dataset and used it to predict the target from the others datasets. The predictive power was measured with the following metrics: R² values to gauge model generalization, error percentages (equation 2) to quantify predictive accuracy, and the percentage of missing taxa in test datasets that was impossible to replace (imputation method returned zero). This simulation was conducted using the data from ‘Group C: cross-studies’, consisting of two subsets, and all taxa were identified by their taxonomy.

The first subset consists of two separate soil experiments with different environmental contexts, still both sequenced identical 16S rRNA variable regions and assessed the pH of the soil. The second subset consists of four human studies, each with different primary research objectives and experimental protocols (sequenced 16S rRNA variable regions, data collection techniques, and material type), nevertheless in all cases patient age was present in the metadata, which was used as a prediction target.

### Comparison of computation complexity and running time

To measure the computational time of CODARFE and compare its performance against other tools, a benchmark was conducted using simulated compositional data organized in two sets of data, as follows: (i) containing one hundred predictors, while the number of samples varied; and (ii) fixed number of samples (n=50), while altering the number of predictors. All experiments were conducted on a Linux Ubuntu 24.04 server with the physical configuration of Xeon E5-2620 v4@2.10GHz and 192GB [2133Mhz DIMM DDR4 Dual Rank] Dell PowerEdge R630. After the fifth day, Coda4Microbiome and Selbal algorithms were prematurely terminated due to excessive time required for each analysis. Only the datasets completed by all tools were considered for comparison.

## Results and Discussion

### Pure correlation outperforms phylogeny for imputation method

Missing predictors is not a common problem in the field of machine learning, as one of the processes in the traditional ML pipeline is predictor extraction [Guyon and Elisseeff, 2006]. In this case the predictors are the taxonomic labels that are used to train the model, but are not necessarily present in the new samples. This heterogeneity in the input data has a deep impact on the prediction outcome, as without the correct information, the model may incorrectly predict the result [Nijman et al., 2022]. Two possibilities were considered to address this problem: the abundance-correlation and the phylogenetic relationship. Although the abundance-correlation may have statistical significance, it might lack biological meaning, as having the same abundance does not necessarily mean two bacteria have the same biological functions [Escalas et al., 2019]. Additionally, there is no guarantee that the most correlated bacteria will exist in the new sample. The phylogenetic relationship approach is based on the idea that if two bacteria are closely related evolutionarily, there is a possibility that they are fulfilling a similar biological role [Gill et al., 2017]. Although the phylogenetic tree is constructed using both new samples and the training dataset, it is possible that the closest evolutionarily related bacterium may not be close enough [Sexton et al., 2017] to have the same function.

As shown in Figure 3, the abundance-correlation method obtained the lowest mean absolute error concerning the percentage of missing data, making it the chosen imputation method for the tool. There may be at least two hypotheses for why the correlation method outperformed the others: (i) The chosen bacterium’s abundance value was closer to the original bacterium’s value, which helped the model correctly predict the target variable. Although the biological explanation suggested the use of techniques based on phylogenetic distance, the abundance values between closely related species can be very different. Since the model does not consider the biological aspect and relies solely on abundance values, pure correlation tends to yield better results; (ii) The second hypothesis is based on the redundancy of biological roles within the microbiome. In this hypothesis, two bacteria with a strong correlation of abundance may possess this similarity precisely because they have redundant roles [Rivett and Bell, 2018].

**Figure 3.**
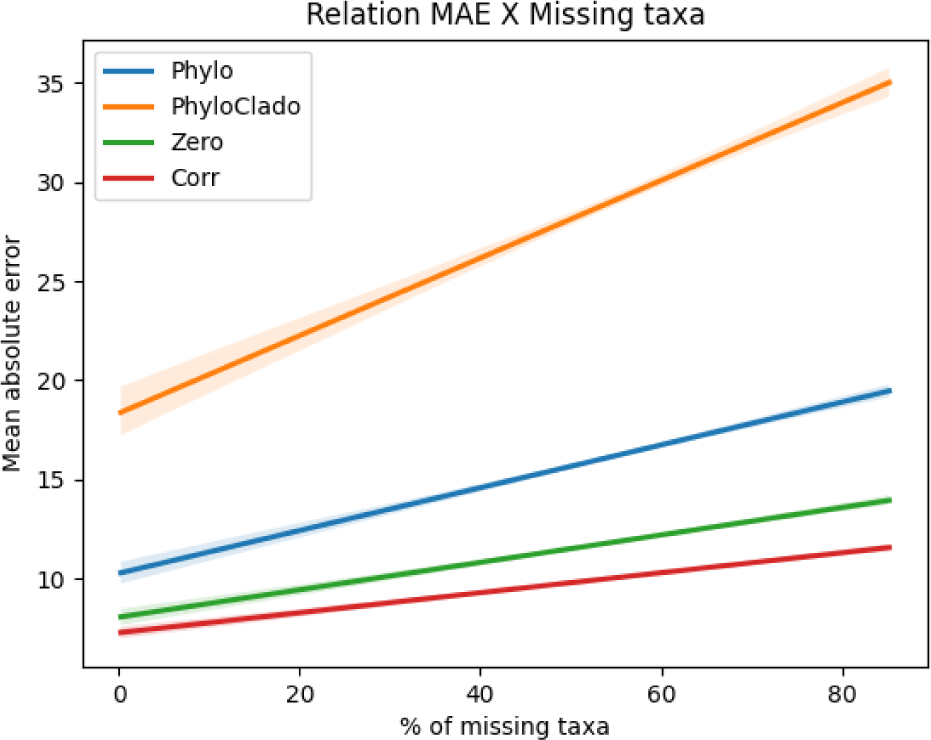
The results of total MAE by percentage of missing taxa for each one of the four tested imputation methods. The database used for testing was the ‘aggregate-stability’ from “Group A: Literature”, with targets ranging from 1.16 to 91.37 units with an average of 21.08 and standard deviation of 17.48. Starting with all taxa, on each step 1% of the total taxa are removed until there is no more taxa present. On each step the MAE is calculated for each one of the four tested imputation methods (Zero, Corr, Phylo and PhyloClade). The MAE is shown in the y-axis and the percentage of missing taxa is presented in the x-axis.

### CODARFE outperformed other tools regarding data generalization

Two groups of datasets were used for benchmarking, 19 soil health metrics (Group A: Literature) and 5 human diseases (Group B: ML Repo). For “Group A: Literature”, CODARFE achieved the best correlation between the generalized and real values for 18 out of the 19 measurements. BRACoD and Coda4Microbiome showed similar results (0.03 units on average) to each other, but the latter failed to generalize the amount of phosphorus (“P”), returning “not a number” (nan) during the process. Selbal had the lowest correlation coefficient among all methods, being unable to generalize the data for 4 out of the 19 measurements, namely “P”, “Mn”, “subsurface hardness”, and “surface hardness” (Figure 4A) (The complete data distribution is shown in Figure S2).

**Figure 4-.**
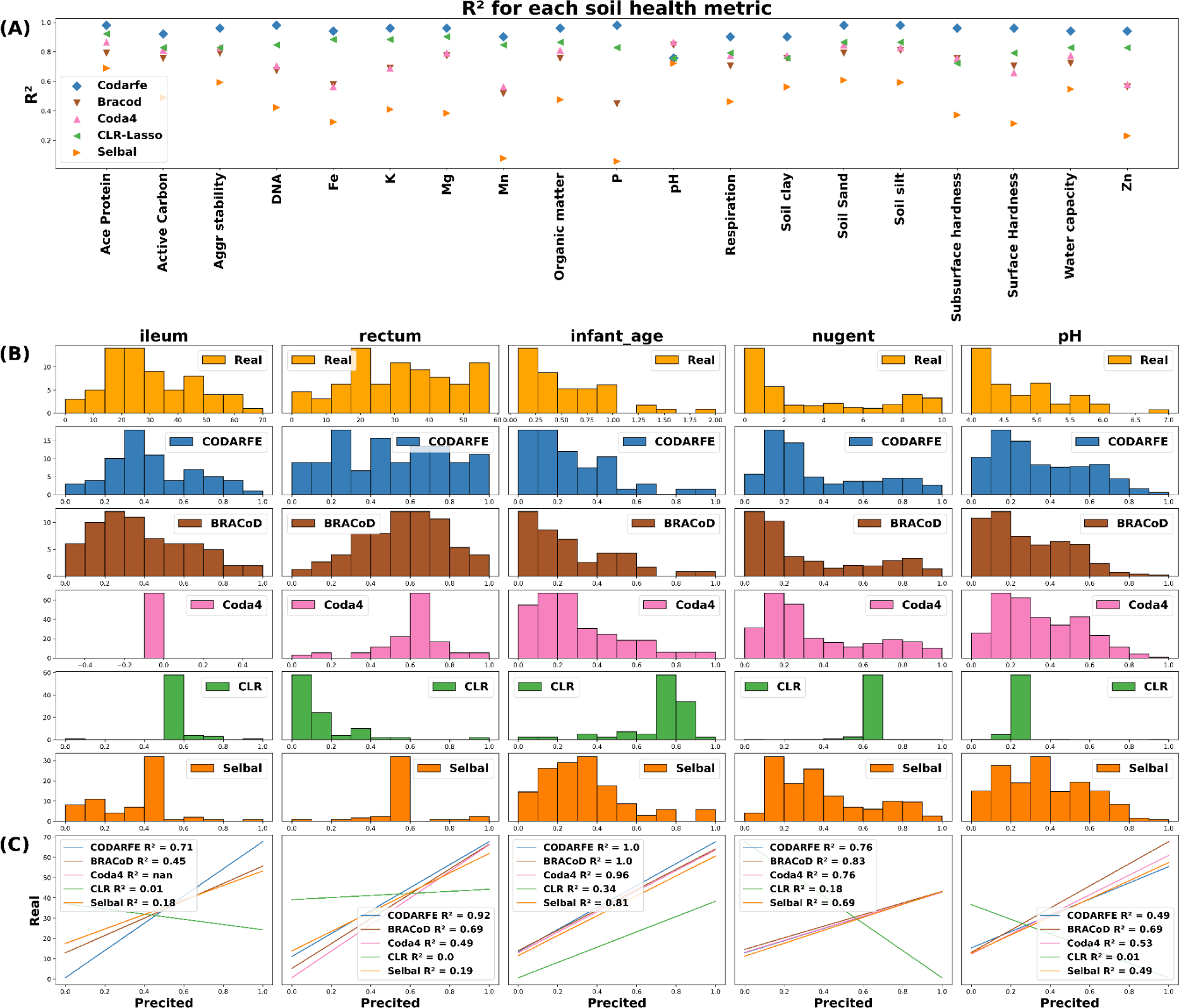
Performance comparison of different machine learning tools for soil and human health prediction. Panel (A): Shows the performance on 19 soil health metrics. CODARFE achieved the best correlation for 18 metrics. BRACoD and Coda4Microbiome had similar performance, but Coda4Microbiome failed to predict phosphorus (P). Selbal had the worst performance. The complete distribution of the data is available in Figure S2.Panel (B): Shows the performance on five human disease datasets through histograms of frequency distribution. A distribution similar to the real one indicates good generalization ability of the model. CODARFE outperformed other tools for pediatric Crohn’s disease. Coda4Microbiome failed to predict ileum samples, and CLR-LASSO failed on all datasets. BRACoD and Coda4Microbiome outperformed CODARFE for vaginal health datasets. Panel (C): Shows the R² values for each method on all datasets. CODARFE achieved the best overall performance except for pH and Nugent score.

For “Group B: ML Repo”, CODARFE outperformed the other tools by at least 0.2 in terms of R² for the pediatric Crohn’s disease dataset. Coda4Microbiome failed to generalize the data for the samples taken from the Ileum, and the CLR-LASSO was unable to generalize any of the datasets. Regarding the vaginal health datasets, BRACoD and Coda4Microbiome outperformed CODARFE (Figure 4), suggesting a complex pattern that is difficult for CODARFE to understand.

Except for pH measurements and Nugent scores, CODARFE outperforms all others in the “learned vs real” correlation. This suggests that the combination of the coefficient shrinkage method with RFE has a higher ability to associate predictors with the target, as previously proposed in [Hamada et al., 2021], where a similar proposal was used and compared with other methods for cervical cancer prediction. Methods primarily based on coefficient shrinkage, such as Coda4microbiome and Selbal, do not test for subsets smaller than the shrinkage suggested. These subsets may hide combinations of predictors that provide better insights. This problem can be mitigated through more exhaustive tests, such as using RFE.

Finally, the lower performance from CODARFE in the vaginal pH and Nugent score datasets may be attributed in part to the way the values were measured. Both datasets had 388 samples, but there were only 11 distinct Nugent score values and 7 pH values. Such a low data variation can affect the generalization performance [He and Garcia, 2009] and make the problems more classifiable rather than prone to regression.

### Simulating prediction in identical studies

The next test involved simulating studies through the hold-out method to simulate predictions. Equation 2 was used to calculate the measure of error (MAPE), which expresses the error as a percentage of each target variable’s mean. Using data from “Group A: Literature” (Figure 5), “surface-hardness” resulted in the lowest error of 0.01% of the target’s mean (target range: 0.001 - 743.126), followed by “Zn” with 3.37% (target range: 0.001 - 23.617) and “soil-texture clay” with 6.21% (target range 0.356 - 39.436). The highest error was found for “respiration” with 47.88% (target range 0.077 - 1.781).

**Figure 5.**
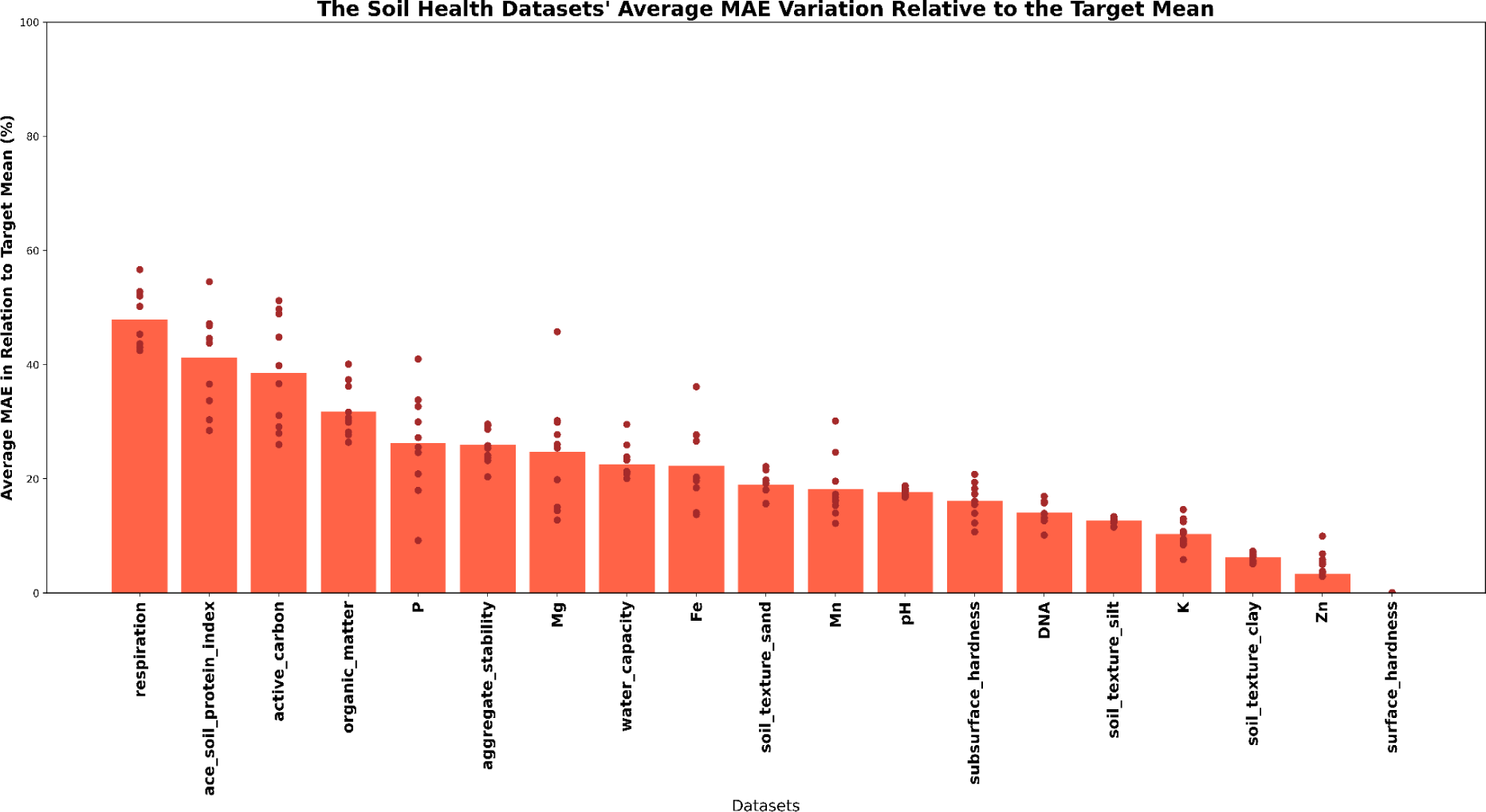
Mean Absolute Percentage Error distribution for soil health and fertility metrics. Individual values across a 10-repetition hold-out validation are represented by the dots, and the average is indicated as the bar. It is possible to see a trend related to the target amplitude, where a narrow distribution has the highest error rate (“respiration”) and a wide distribution has the lowest error (“surface_hardness”).

One hypothesis for this arising pattern is related to the data distribution. The “respiration” metric had the narrowest distribution of all the metrics (Figure S3), varying only 1.7 units from minimum to maximum with a standard deviation of 0.23 units, making any deviation from the prediction to the true value to result in a high error. In contrast, the “surface_hardness” had one of the most widely distributed distributions, ranging from 0 to over 700 units with a standard deviation of 105 units. On top of that, the narrow amplitude has high peaks of density, which may create a bias in the model reflecting in even higher errors when the true value is outside the high density range [Gyori et al., 2022]. On the other hand, most of the results obtained were superior (in terms of R²) than the reported in the original article [Wilhelm et al., 2022], supporting the high predictive power of CODARFE in most of the scenarios.

### CODARFE predictor selection method surpasses state-of-art in terms of proportions of correctly selected

For a more in-depth analysis of the human microbiome data contained within the “Group B: ML Repo”, we tried to verify the relevance of the selected taxa by the different tools. The three publications that were used are each briefly detailed here, along with a description of the taxa that were discovered to be connected to the interest variable.

Yatsunenko et al. (2012) conducted a study utilizing fecal samples from diverse populations, including healthy Amerindians from the Amazon and Venezuela, rural communities in Malawi, and metropolitan areas in the United States. Their aim was to characterize postnatal developmental stages and their correlation with environmental factors. Although the study did not explicitly seek direct associations between the gut microbiome bacteria and postnatal age, Bifidobacteruim was pointed out by the authors as a potential biomarker for infant age. As shown in green in the bar plots (Figure 6), only CODARFE and BRACoD selected Bifidobacterium species, with CODARFE selecting 41 times less false-positives than BRACoD. The other selected taxa (red) are not supported by the original article.

**Figure 6.**
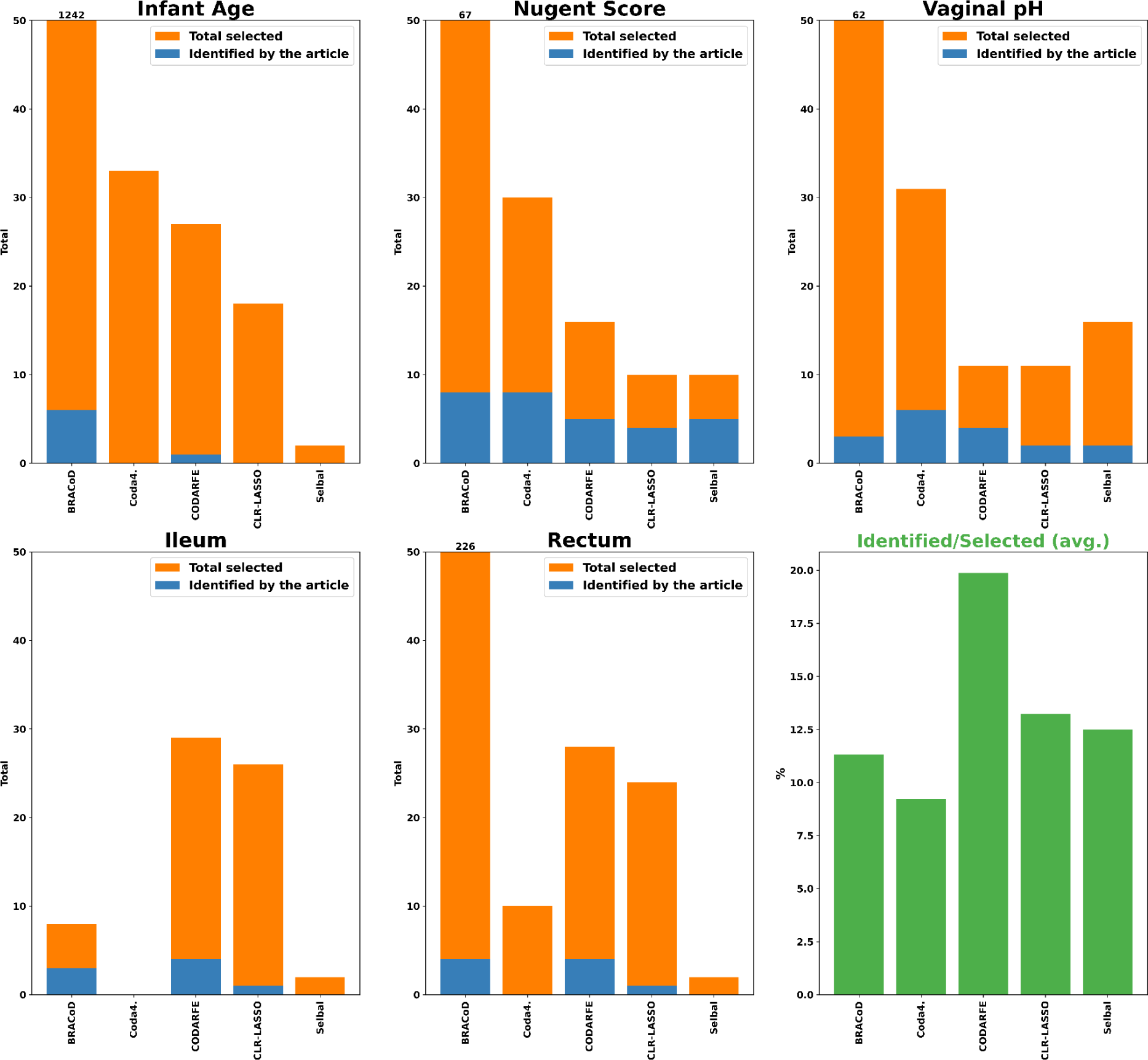
The figure illustrates the performance of all five tools for selecting relevant taxa identified in three different studies (with 5 different target variables) on the human microbiome. The bars show the number of taxa correctly selected by each tool (blue) and the number of taxa not supported by the original study (orange) for each dataset. CODARFE generally selected more taxa identified in the original studies compared to other tools, and also had a lower proportion of false positives (taxa not supported by the studies) compared to other tools as illustrated by the last panel (green).

Ravel et al. (2010) characterized bacteria present in vaginal samples from four ethnic groups (White, Black, Hispanic, and Asian) to ascertain vaginal pH and Nugent scores. The following phylotypes were associate with high Nugent scores *Aerococcus, Anaeroglobus, Anaerotruncus, Atopobium, Coriobacteriaceae 2, Dialister, Eggerthella, Gardnerella, Gemella, Megasphaera, Mobiluncus, Parvimonas, Peptoiphilus, Prevotella, Porphyomonas, Prevotellaceae1, Prevotellaceae 2, Ruminococcaceae*, and *Snethia*. Conversely, phylotypes of *Lactobacillus* were associated with a low Nugent. From the 20 highlighted phylotypes, BRACoD correctly selected a total of 8 different phylotypes being them: *Anaeroglobus, Prevotella, Lactobacillus, Gemella, Atopobium, Aerococcus, Gardnerella,* and *Dialister*; Coda4Microbiome correctly selected 8 phylotypes being them: *Mobiluncus, Prevotella, Lactobacillus, Gemella, Atopobium, Aerococcus, Gardnerella,* and *Dialister*; CODARFE correctly selected 5 phylotypes being them: *Prevotella, Lactobacillus, Atopobium, Gardnerella,* and *Dialister*; CLR-LASSO correctly selected a total of 4 phylotypes being them: *Prevotella, Lactobacillus, Gemella,* and *Atopobium*; and Selbal selected correctly 5 phylotypes being them: *Prevotella, Lactobacillus, Atopobium, Gardnerella,* and *Dialister*. For vaginal pH, only the presence or absence of *Lactobacillus* was reported to be relevant, with its presence being inversely proportional to the pH measurement. BRACoD selected 3 species; Coda4Microbiome selected 6 species; CODARFE selected 4 species; CLR-LASSO selected 2 species; and finally Selbal selected 2 species. It is worth noting that the proportions of correctly selected and total selected vary drastically between tools (Figure 6).

The work developed by Gevers et al. (2014) characterized bacteria present in ileum and rectal samples to understand recent cases of Crohn’s disease in children (PCDA). The authors identified 21 species with a high correlation to PCDA: *Escherichia coli, Fusobacterium nucleatum, Haemophilus parainfluenzae, Veillonella parvula, Eikenella corrodens, Gemella moribillum, Bacteroides vulgatus, Bacteroides caccae, Bifidobacterium bifidum, Bifidobacterium longum, Bifidobacterium adolescentis, Bifidobacterium dentum, Blautia hansenii, Ruminococcus gnavus, Clostridium nexile, Faecalibacterium prausnitzii, Ruminoccus torques, Clostridium bolteae, Eubacterium rectale, Roseburia intestinalis,* and *Coprococcus comes*. The results in figure 6 are shown for two separated datasets (ileum and rectum), but all species are valid for both datasets. BRACoD correctly selected *Haemophilus parainfluenzae, Veillonella parvula, and Bacteroides vulgatus* for the Ileum dataset and *Fusobacterium nucleatum Eikenella corrodens, Bifidobacterium bifidum,* and *Bifidobacterium longum* for rectum dataset; As previously described, Coda4Microbiome was unable to finish its process for the Ileum dataset, and did not select any of the 21 species pointed out by the article in the rectum dataset; CODARFE correctly selected *Fusobacterium nucleatum, Bacteroides vulgatus, Faecalibacterium prausnitzii,* and *Roseburia intestinalis* for ileum dataset and *Haemophilus parainfluenzae, Bacteroides vulgatus, Bifidobacterium longum,* and *Eubacterium rectale* for rectum dataset; CLR-LASSO selected only *Veillonella parvula* for ileum and rectum; and Selbal did not select any of the 21 species.

In these three cases, CODARFE finds more species highlighted in the original articles compared to other tools (green bar chart in figure 6), suggesting that it has lower False Discovery Rate (FDR) than the other tools on average. We believe that CODARFE was able to achieve these results due to the combination of metrics with the RFE [Guyon and Elisseeff, 2003]. It has two strong indications of a high quantity of predictors, being the R^2^ adjusted and the BIC metrics [Schwarz, 1978a,b]. Both these metrics account as a penalization metric for the number of predictors selected, giving high scores as the number of predictors decrease. In addition, we combined them with the ρ-value of the F-test in the final predictors score. This value basically indicates how beneficial the inclusion of a predictor is to the model (it can be thought of as a value representing “how much these predictors explain the target”). Since these three metrics are weighted and summed together, the final score will indicate “how worthwhile it is to keep all these predictors in relation to how much they explain”. This value is weighted with an “overfitting value” and used to evaluate all (at most) 100 predictors sets during the RFE step, increasing the chance of finding a subset of predictors with optimal scores that are not explored by other methods (such as shrinkage used by Coda4Microbiome) [Guyon et al., 2002].

### Cross-studies

We separated “Group C: cross-studies” into two groups, one similar to each other and one highly different, in order to better examine the effects of studies that were similar and studies that were quite different in many areas, such as subject of investigation, region of sequencing, data collection, external effects, and others (the differences between each dataset are indicated in Tables 2 and 3). To assess CODARFE’s robustness we used the R² values to judge model generalization (Figures 7C and 8B), mean absolute percentage error (MAPE) to quantify predictive accuracy (Figure 7B and 8C), and the percentage of missing taxa in test datasets (imputation method returned zero) to verify the effect of missing taxa in the final prediction (Figure 8D).

**Figure 7.**
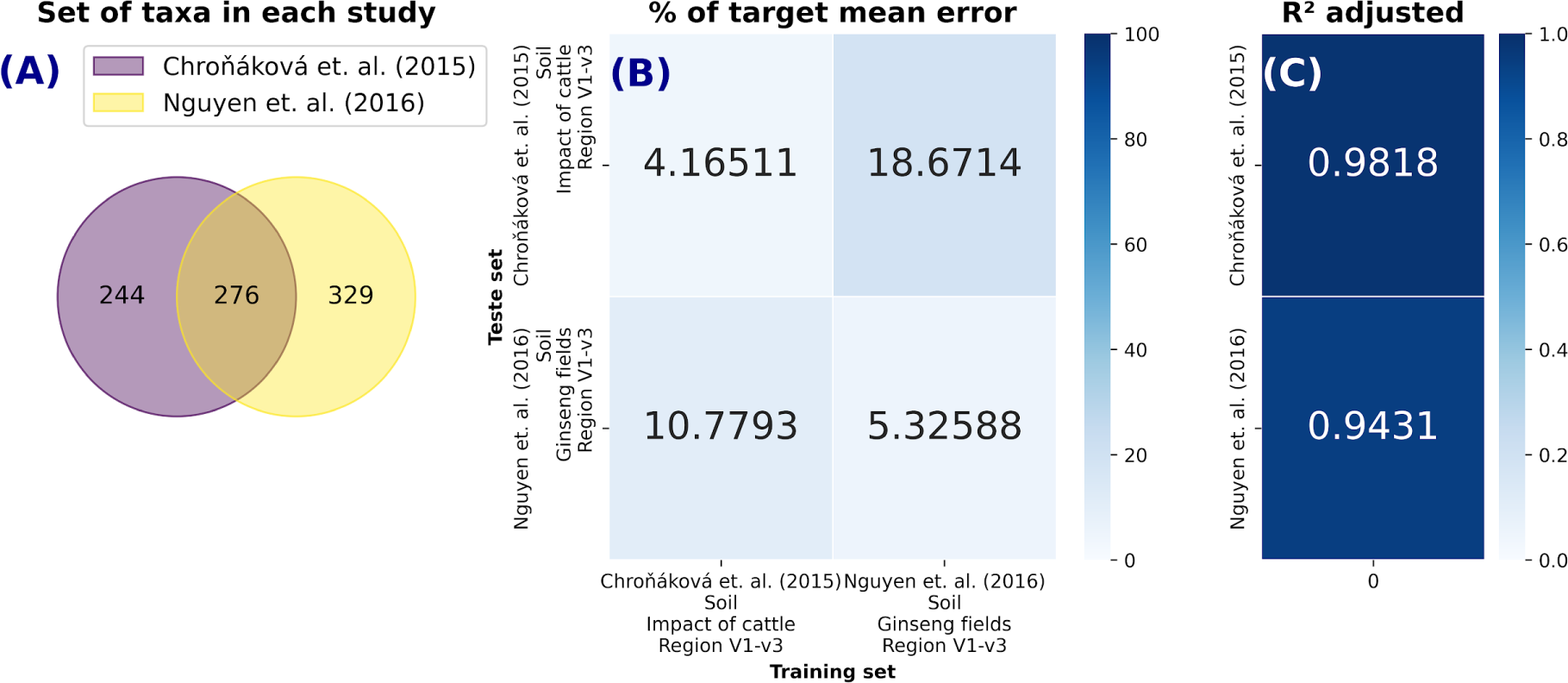
Cross-studies prediction for tow soil datasets measuring pH. The number of identical predictors in each study are shown in (A). The percentage of mean absolute error in relation to the average of the target variable when training CODARFE with data from one work and predicting another is shown in (B). The coefficient of determination (*R*^2^) obtained in the generalization of the CODARFE model is shown in (C).

**Figure. 8.**
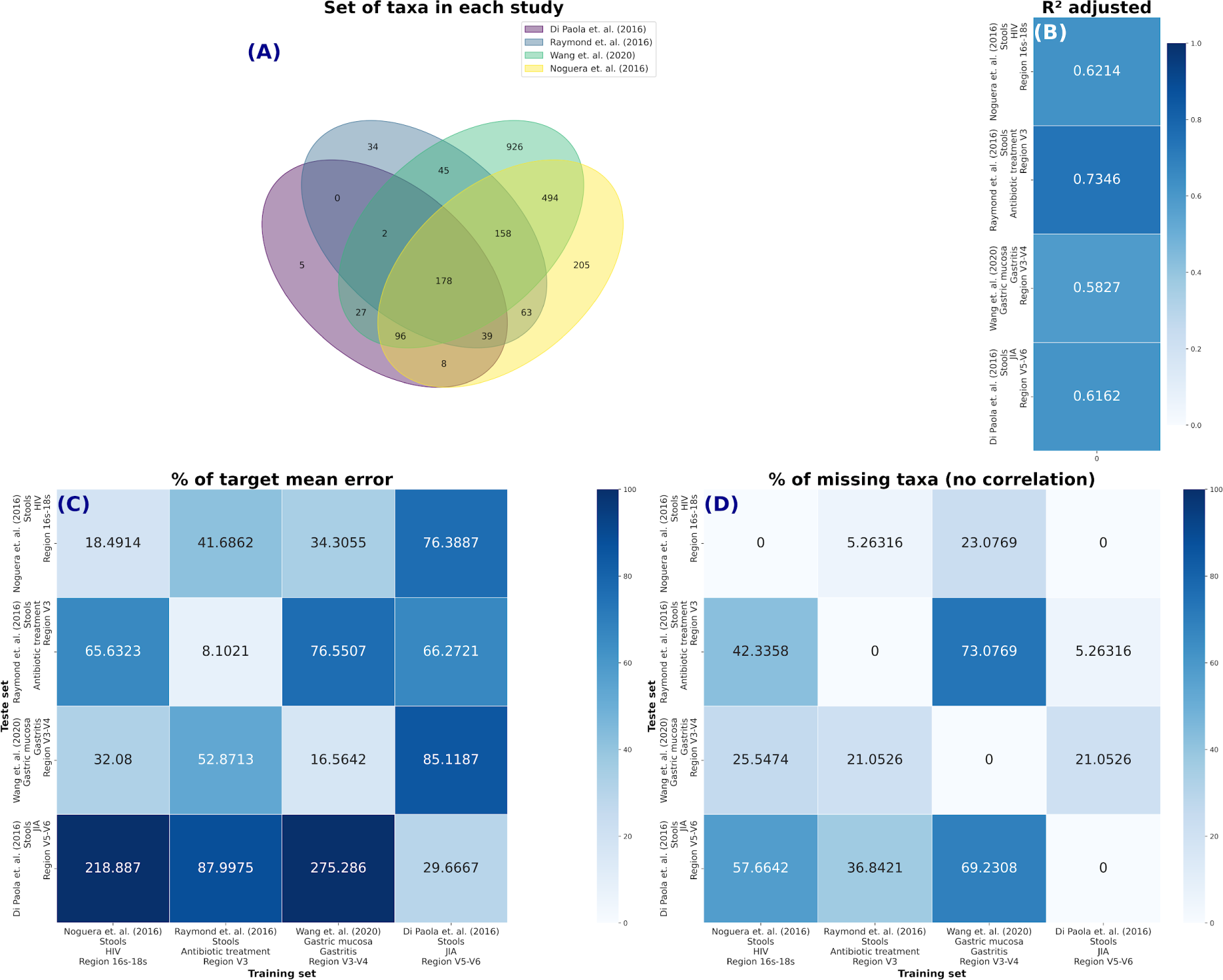
Cross-studies prediction for four very distinct human datasets measuring the human age. The number of identical predictors (taxa) in each study is shown in (A); The coefficient of determination (*R*^2^) obtained in the generalization of the CODARFE model is shown in (B); The percentage of mean absolute error in relation to the average of the target variable when training CODARFE with data from one work and predicting another is shown in (C); The percentage of predictors in the prediction dataset that were unable to be replaced by the imputation method (the imputation method returned zero) is shown in (D).

**Table 2.** Details of each dataset used in the soil cross-studies analysis.

|  | Chroňáková et. al. (2015) | Nguyen et. al. (2016) |
| --- | --- | --- |
| **Sample source** | Soil | Soil |
| **Aim of the study** | Investigation of prokaryotic diversity in soil impacted by outdoor cattle husbandry | Investigation of bacterial communities in the soil of ginseng fields |
| **Location** | Czech republic | South Korea |
| **Year of collection** | 2011 | 2012 |
| **Sequencing platform** | Illumina | Illumina |
| **Sequenced region** | V1-V3 | V1-V3 |
| **MGnify study identifier** | MGYS00000916 | MGYS00001160 |

**Table 3.** Table exhibiting pertinent data from each research paper used in the human cross-studies analysis.

|  | Di Paola et. al.<br>(2016) | Raymond et. al.<br>(2016) | Wang et. al.<br>(2020) | Noguera et. al.<br>(2016) |
| --- | --- | --- | --- | --- |
| **Sample** | Large intestine | Large intestine | Digestive system | Large intestine |
| **Sample type** | Feces | Feces | Gastric mucosa | Feces |
| **Aim of the Study** | Characterization of gut microbiota in Juvenile Idiopathic arthritis | Study of the impact of antibiotics on the gut microbiome | Microbiota in patients with gastritis, intestinal metaplasia, early gastric cancer and advanced gastric cancer | Understanding the impact of HIV on the microbiome |
| **Platform** | Illumina | Illumina | Illumina | Illumina |
| **Sequenced region** | V5-V6 | V3 | V3-V4 | 16s/18s rRNA |
| **Host** | Children | Adults between 21 and 35 years old | Adults | Adults |
| **MGnify study identifier** | MGYS00000580 | MGYS00001175 | MGYS00001188 | MGYS00001255 |

In the first subset, Chrǒnáková et al. (2015) evaluated the response of soil bacterial communities to changes associated with outdoor cattle overwintering, while Nguyen et. al. (2016) analyzed how bacterial diversity and community structure in Korean ginseng fields were altered by cultivation time. Both datasets achieved high R² (Figure 7C) which indicates a high predictive power. For the Chrǒnáková et. al. (2015), the error percentages were observed to be 4.17% and 10.78% (Figure 7B) when tested on itself and the Nguyen et. al. (2016) dataset, respectively. Similarly, the Nguyen et. al. (2016) exhibited error percentages of 5.33% and 18.67% (Figure 7B) when tested on the Chrǒnáková et. al. (2015) dataset and itself, respectively.

Both the Chrǒnáková et. al. and Nguyen et. al. (2016) datasets were generated from the 16S rRNA V1-V3 region (Table 2), and all the predictors (taxa) selected by CODARFE were present in both studies (in the intersection, Figure 7A), meaning that no imputation was needed, and reinforcing the idea that pH related taxa are similar for various types of soils [Bååth, 1996]. This will have helped with the low error even considering the agent of the study effect is entirely different, and the soil samples are at least 8,000 kilometers apart. The sequenced region directly influences the assigned taxonomy [Guo et al., 2013], which is then used as a predictor by the tool, since, if different taxonomies are assigned to the same predictor, they will be considered as different predictors. With these results, a key factor influencing the observed prediction errors may be the choice of amplified region, since both the Chrǒnáková. et. al. (2015) and Nguyen et. al. (2016) datasets were generated from the V1-V3 region (Table 2). This factor is better observed in the second subset.

In the second subset, four completely different studies were used (Table 3). Raymond et al. (2016) model demonstrates the lowest error when tested on itself (8.10%) (Figure 8C) and exhibits the lowest error on average (53.41%) when predicting others datasets. Conversely, the model by Di Paola et al. (2016) yields the highest average error (80.11%) (Figure 8C).

The best results yielded an error of 32% of the mean (Figure 8C), indicating low predictive power between studies. A reduced R^2^ in these studies suggested that the microbiota may be more influenced by the study object than by age. Furthermore, although the case with the best error (32%) has a 25% missing taxa rate (Figure 8D), it is not only the absence of taxa that affects the error rate, as shown by the model trained with Di Paola et. al. (2015), which was unable to predict the age accurately in other works, despite containing most taxa. This may be partially explained by the fact that this work had the lowest fitting power (R2) (Figure 8B) among all, leading us to believe that the error may be a combination of fitting power with missing taxa and sequencing region.

The influence of missing taxa becomes evident when comparing the models trained on datasets with fewer missing taxa to the highest ones. This trend can be observed with the Wang et al. (2020) model trained on the dataset with the lowest missing taxa percentage (0%) also demonstrating the lowest average error across test datasets (Figure 8C and D). This trend is reaffirmed by the challenges faced by the Wang et. al. (2020) and Noguera et. al. (2016) models in predicting the data of Di Paola et. al. (2016) highlight the effects of missing taxa and sequencing regions in the model effectiveness. The percentage of missing taxa in the Di Paola et. al. (2016) dataset (Figure 8D) is substantially higher compared to the training datasets. The Wang et. al. (2020) model, trained on V3–V4 region, and the Noguera et. al. (2016) model, trained on 16S/18S rRNA (Table 3), might not have adequately captured predictors relevant to the V5-V6 region utilized by the Di Paola et. al. (2016) dataset (Figure 8A and D). This may suggest a critical relationship between missing taxa, and predictive accuracy.

Further examining the sequencing regions used to generate the data, it again suggested that different regions can impact the predictive performance. For instance, the Noguera et al. (2016) model, trained on 16S/18S rRNA data (Table 3), exhibits relatively strong cross-dataset performance, while the Raymond et al. (2016) model, trained on the V3 region (Figure 8C) showcases the highest R² value and lowest error on average. Another indication that the sequenced region is an essential factor in the tool’s proper functioning comes from the cross-study analyses of Di Paola et. al. (2016) and Wang et. al. (2020). These two datasets did not share the 16S sequenced region and resulted in the highest error rates. Conversely, the models created from Wang et. al. (2020) and Raymond et. al. (2016), both sequencing the V3 region, when used to predict Noguera et. al. (2016) study, which sequenced the complete 16S, yielded errors of 34% and 41%, respectively, which are relatively low compared to its other case with 275% error.

Based on the results obtained in both cross-study analyses, we elaborated three hypotheses regarding CODARFE feasibility and limitations:

i. The main factor determining predictive power is the number of predictors that could not be replaced by the suggested imputation method, as observed in the models trained with Wang et. al. (2020) or Noguera et. al. (2016) datasets and tested in Di Paola et. al. (2016) and Raymond et. al. (2016) datasets. These models had the highest associated error and the highest percentage of missing predictors. Considering such a limitation, it is likely that the sequenced region needs to be the same for better results since the sequenced region directly affects the assigned taxonomy.
ii. The samples from which the sequences are extracted should be similar, as the microbiome is closely linked to the environment. Thus, samples from different types (e.g.: Feces and gastric mucosa) also negatively impact the model’s predictive ability, as seen in the models trained with Wang et. al. (2020) dataset.
iii. Lastly, the “objective of the study” for new samples should be similar to that used for training. For example, the model generated from Di Paola et. al. (2016), which focused on children, could not accurately predict ages in other studies, indicating that the taxa associated with infant age differ from those associated with adult age, supporting the idea discussed in [Yatsunenko et al., 2012].

### CODARFE has linear runtime growth with respect to the number of predictors and samples

Figure 9 shows the computational time used to perform the analysis for each tool. Regarding the number of samples, BRACOD demonstrated a considerable increase in runtime correlated with increased sample sizes, Coda4Microbiome and CLR-LASSO exhibited the lowest runtimes. Selbal had a moderate upswing in runtime, followed closely by CODARFE, but none surpasses 400s (6 minutes) per database.

**Figure 9.**
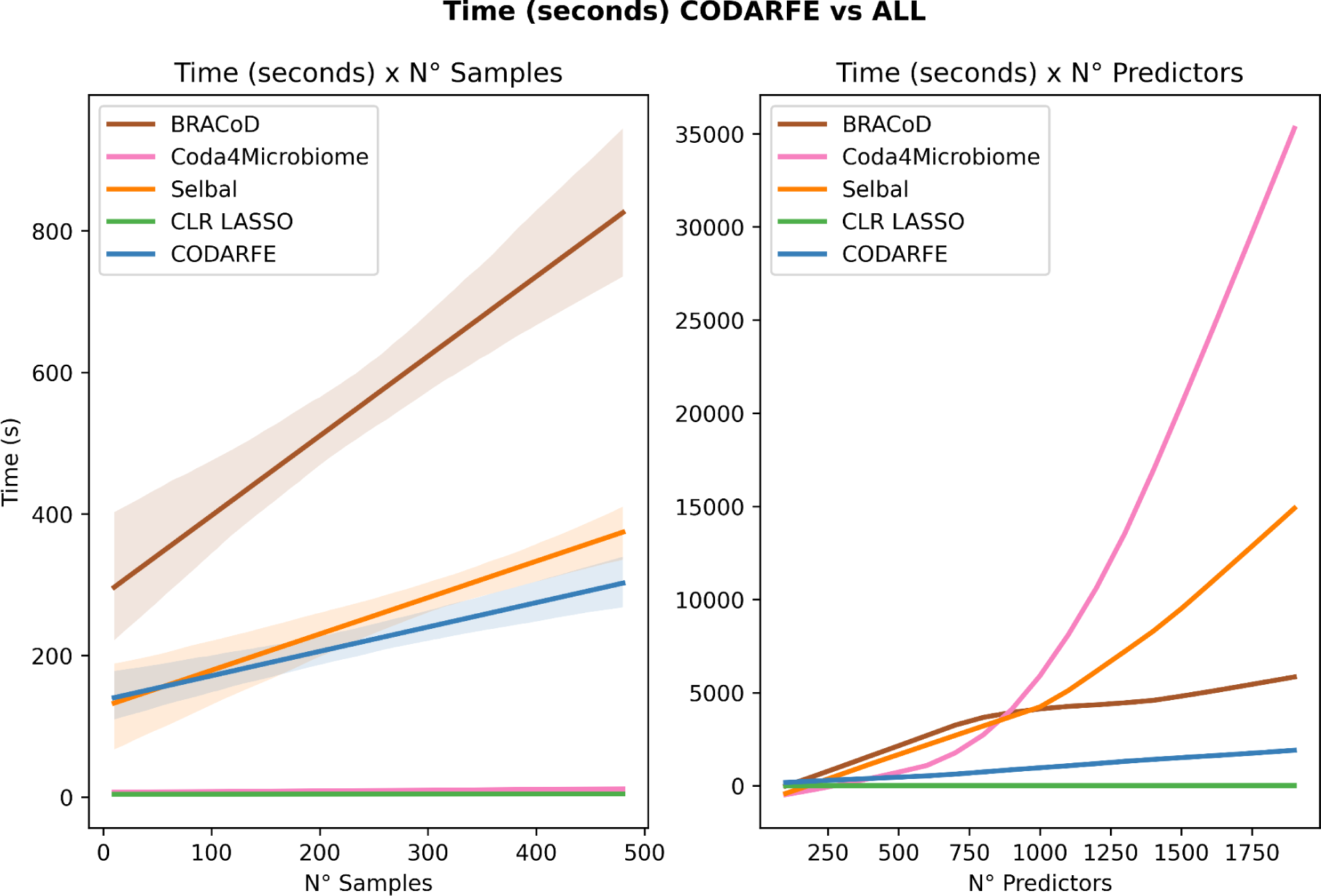
Time consumption comparison between tools regarding number of samples (rows) and number of predictors (columns) in the training dataset. The graph on the left depicts the time in seconds required by each tool to finish its process as the number of samples in the database increases. The graph on the right depicts the time in seconds required by each tool to finish its process as the number of predictors in the database increases.

Regarding the number of predictors, the data transformation used by Coda4microbiome and Selbal tools consumes quadratic time, increasing its computational costs and limiting its usage of datasets with high numbers of predictors (e.g.: >1500 predictors), what can become a problem since the growth of the number of predictors (columns) is being driven by the development of more data-generating sequencing techniques [Quince et al., 2017]. The BRACoD’s execution time significantly increases with the number of predictors. Furthermore, there is a limitation of 300 predictors suggested by the authors together with the “solution” of selecting just the top 300 most abundant taxa in the dataset. This may raise a fundamental issue in microbiome, which is that the most abundant taxa may not be the most relevant to the target variable [Woodcock et al., 2006].

While CLR-LASSO has the lowest time in both cases, interpreting its results can be challenging due to the applied transformation and the effect of the mean of non-selected variables. Since this transformation is used in the whole dataset and just a few predictors are selected, the analysis of the method suffers effects from the unselected predictors. Furthermore, the user’s choice of the value of lambda directly affects the number of selected predictors, and it lacks generalization capability as evidenced by previous trials.

Finally, CODARFE remain computationally viable even when a high number of predictors (e.g., > 1500) are utilized, making it the best choice for largest datasets.

## Conclusion

We present CODARFE, a powerful tool capable of selecting sparse compositional predictors and associating them with continuous environmental variables, which in turn can be used to predict these variables in new samples. CODARFE’s performance was benchmarked against four other tools and exhibited superior results in 21 out of 24 tested datasets, with over 0.1 units stronger in terms of R² on average, and 17 times faster than the most recent published tools, particularly when the number of predictors surpassed 1500. For human-related data, CODARFE achieved the highest number of identified taxa linked to the target variable that were also identified in the publication describing the study. Furthermore, we demonstrated CODARFE’s predictive power in cross-study analyses, where a model trained on data from one project could be successfully used to predict data from another project, with a maximum error rate of 18% in relation to the target variable mean, as long as the sequence region and sample type are the same. These results indicate CODARFE’s potential for generalization and robustness across multiple studies, even under different experimental settings. Furthermore, the CODARFE uniqueness lies in its ability to predict the target variable in new samples, due to its imputation method for missing predictors. This feature allows for predicting environmental variables in previously unexplored contexts, providing researchers with a reliable and versatile tool. Furthermore, while not demonstrated here, we believe that CODARFE is applicable to data other than taxa, such as protein functions and metabolic pathways, since the mathematical constraints are the same for all of them.

## Data Availability

All tool versions can be found at https://github.com/alerpaschoal/CODARFE, and https://doi.org/10.5281/zenodo.12751711 contains the data and code used to create all of the analyses and plots.

## Acknowledgments

Authors would like to thank the “Postgraduate Associate Program in Bioinformatics (PPGAB)” between UTFPR and UFPR and the Group of Pattern Recognition and Bioinformatics also from UTFPR for the mentoring. M.C.B acknowledges a PhD fellowship from SUPERBAC Biotechnology Solutions and mentoring. ARP was supported by Fundação Araucária - NAPI Bioinformática (# 66.2021) and by CNPq (# 440412/2022-6). RDF is supported by EMBL.

## References

J. Aitchison. The statistical analysis of compositional data. Journal of the Royal Statistical Society: Series B (Methodological), 44(2):139–160, 1982.

E. Bååth. Adaptation of soil bacterial communities to prevailing ph in different soils. FEMS Microbiology Ecology, 19(4):227–237, 1996.

L. Breiman. Random forests. Machine learning, 45:5–32, 2001.

M. L. Calle, M. Pujolassos, and A. Susin. coda4microbiome: compositional data analysis for microbiome cross-sectional and longitudinal studies. BMC bioinformatics, 24(1):82, 2023.

V. Chandrasekhar. Disease2Vec: a method of determining disease from gut microbiome using neural embeddings. PhD thesis, Harvard University, 2020.

C.-C. Chang and C.-J. Lin. Libsvm: a library for support vector machines. ACM transactions on intelligent systems and technology (TIST), 2(3):1–27, 2011.

A. Chrłáková, B. Schloter-Hai, V. Radl, D. Endesfelder, C. Quince, D. Elhottová, M. Šimek, and M. Schloter. Response of archaeal and bacterial soil communities to changes associated with outdoor cattle overwintering. PLoS One, 10(8):e0135627, 2015.

G. L. Damhorst, M. W. Adelman, M. H. Woodworth, and C. S. Kraft. Current capabilities of gut microbiome–based diagnostics and the promise of clinical application. The Journal of Infectious Diseases, 223(Supplement 3):S270–S275, 2021.

X. Dang, H. Peng, X. Wang, and H. Zhang. Theil-sen estimators in a multiple linear regression model. Olemiss Edu, 2008.

M. Di Paola, D. Cavalieri, D. Albanese, M. Sordo, M. Pindo, C. Donati, I. Pagnini, T. Giani, G. Simonini, A. Paladini, et al. Alteration of fecal microbiota profiles in juvenile idiopathic arthritis. associations with hla-b27 allele and disease status. Frontiers in microbiology, 7:1703, 2016.

A. Escalas, L. Hale, J. W. Voordeckers, Y. Yang, M. K. Firestone, L. Alvarez-Cohen, and J. Zhou. Microbial functional diversity: From concepts to applications. Ecology and Evolution, 9(20):12000–12016, 2019.

D. Gevers, S. Kugathasan, L. A. Denson, Y. Vázquez-Baeza, W. Van Treuren, B. Ren, E. Schwager, D. Knights, S. J. Song, M. Yassour, et al. The treatment-naive microbiome in new-onset crohn’s disease. Cell host & microbe, 15(3):382–392, 2014.

R. B. Ghannam and S. M. Techtmann. Machine learning applications in microbial ecology, human microbiome studies, and environmental monitoring. Computational and Structural Biotechnology Journal, 19:1092–1107, 2021.

J. A. Gilbert, M. J. Blaser, J. G. Caporaso, J. K. Jansson, S. V. Lynch, and R. Knight. Current understanding of the human microbiome. Nature medicine, 24(4):392–400, 2018.

A. S. Gill, A. Lee, and K. L. McGuire. Phylogenetic and functional diversity of total (dna) and expressed (rna) bacterial communities in urban green infrastructure bioswale soils. Applied and Environmental Microbiology, 83(16):e00287–17, 2017.

G. B. Gloor, J. M. Macklaim, V. Pawlowsky-Glahn, and J. J. Egozcue. Microbiome datasets are compositional: and this is not optional. Frontiers in microbiology, 8:2224, 2017.

F. Guo, F. Ju, L. Cai, and T. Zhang. Taxonomic precision of different hypervariable regions of 16s rrna gene and annotation methods for functional bacterial groups in biological wastewater treatment. PloS one, 8(10):e76185, 2013.

I. Guyon and A. Elisseeff. An introduction to variable and feature selection. Journal of machine learning research, 3(Mar):1157–1182, 2003.

I. Guyon and A. Elisseeff. An introduction to feature extraction. In Feature extraction: foundations and applications, pages 1–25. Springer, 2006.

I. Guyon, J. Weston, S. Barnhill, and V. Vapnik. Gene selection for cancer classification using support vector machines. Machine learning, 46:389–422, 2002.

N. G. Gyori, M. Palombo, C. A. Clark, H. Zhang, and D. C. Alexander. Training data distribution significantly impacts the estimation of tissue microstructure with machine learning. Magnetic resonance in medicine, 87(2):932–947, 2022.

M. Hamada, J. J. Tanimu, M. Hassan, H. A. Kakudi, and P. Robert. Evaluation of recursive feature elimination and lasso regularization-based optimized feature selection approaches for cervical cancer prediction. In 2021 IEEE 14th International Symposium on Embedded Multicore/Many-core Systems-on-Chip (MCSoC), pages 333–339. IEEE, 2021.

H. He and E. A. Garcia. Learning from imbalanced data. IEEE Transactions on Knowledge and Data Engineering, 21(9):1263–1284, 2009. doi: 10.1109/TKDE.2008.239.

S. Huang, E. Ailer, N. Kilbertus, and N. Pfister. Supervised learning and model analysis with compositional data. PLOS Computational Biology, 19(6):e1011240, 2023.

C. L. Johnson and J. Versalovic. The human microbiome and its potential importance to pediatrics. Pediatrics, 129(5):950–960, 2012.

M. Kuhn. Building predictive models in r using the caret package. Journal of statistical software, 28:1–26, 2008.

Z. D. Kurtz, C. L. Müller, E. R. Miraldi, D. R. Littman, M. J. Blaser, and R. A. Bonneau. Sparse and compositionally robust inference of microbial ecological networks. PLoS computational biology, 11(5):e1004226, 2015.

A. Lavelle and C. Hill. Gut microbiome in health and disease: emerging diagnostic opportunities. Gastroenterology Clinics, 48(2):221–235, 2019.

P. Legendre and E. D. Gallagher. Ecologically meaningful transformations for ordination of species data. Oecologia, 129:271–280, 2001.

H. Lin and S. D. Peddada. Analysis of microbial compositions: a review of normalization and differential abundance analysis. NPJ biofilms and microbiomes, 6(1):60, 2020.

K. C. Lutz, S. Jiang, M. L. Neugent, N. J. De Nisco, X. Zhan, and Q. Li. A survey of statistical methods for microbiome data analysis. Frontiers in Applied Mathematics and Statistics, 8:884810, 2022.

K. W. Ng, G.-L. Tian, and M.-L. Tang. Dirichlet and related distributions: Theory, methods and applications. 2011.

N.-L. Nguyen, Y.-J. Kim, V.-A. Hoang, S. Subramaniyam, J.-P. Kang, C. H. Kang, and D.-C. Yang. Bacterial diversity and community structure in korean ginseng field soil are shifted by cultivation time. PloS one, 11(5):e0155055, 2016.

S. Nijman, A. Leeuwenberg, I. Beekers, I. Verkouter, J. Jacobs, M. Bots, F. Asselbergs, K. Moons, and T. Debray. Missing data is poorly handled and reported in prediction model studies using machine learning: a literature review. Journal of clinical epidemiology, 142:218–229, 2022.

M. Noguera-Julian, M. Rocafort, Y. Guillén, J. Rivera, M. Casadellà, P. Nowak, F. Hildebrand, G. Zeller, M. Parera, R. Bellido, et al. Gut microbiota linked to sexual preference and hiv infection. EBioMedicine, 5:135–146, 2016.

N. Novikova, P. De Boever, S. Poddubko, E. Deshevaya, N. Polikarpov, N. Rakova, I. Coninx, and M. Mergeay. Survey of environmental biocontamination on board the international space station. Research in microbiology, 157(1):5–12, 2006.

A. B. Owen. A robust hybrid of lasso and ridge regression. Contemporary Mathematics, 443(7):59–72, 2007.

C. Quince, A. W. Walker, J. T. Simpson, N. J. Loman, and N. Segata. Shotgun metagenomics, from sampling to analysis. Nature biotechnology, 35(9):833–844, 2017.

J. Ravel, P. Gajer, Z. Abdo, G. M. Schneider, S. S. Koenig, S. L. McCulle, S. Karlebach, R. Gorle, J. Russell, C. O. Tacket, et al. Vaginal microbiome of reproductive-age women. Proceedings of the National Academy of Sciences, 108(supplement 1):4680–4687, 2011.

F. Raymond, A. A. Ouameur, M. Déraspe, N. Iqbal, H. Gingras, B. Dridi, P. Leprohon, P.-L. Plante, R. Giroux, È. Bérubé, et al. The initial state of the human gut microbiome determines its reshaping by antibiotics. The ISME journal, 10(3):707–720, 2016.

L. Richardson, B. Allen, G. Baldi, M. Beracochea, M. L. Bileschi, T. Burdett, J. Burgin, J. Caballero-Pérez, G. Cochrane, L. J. Colwell, et al. Mgnify: the microbiome sequence data analysis resource in 2023. Nucleic Acids Research, 51(D1):D753–D759, 2023.

J. Rivera-Pinto, J. J. Egozcue, V. Pawlowsky-Glahn, R. Paredes, M. Noguera-Julian, and M. L. Calle. Balances: a new perspective for microbiome analysis. MSystems, 3(4):10–1128, 2018.

D. W. Rivett and T. Bell. Abundance determines the functional role of bacterial phylotypes in complex communities. Nature microbiology, 3(7):767–772, 2018.

G. Schwarz. Estimating the Dimension of a Model. The Annals of Statistics, 6(2):461–464, 1978a. doi: 10.1214/aos/1176344136. URL https://doi.org/10.1214/aos/1176344136.

G. Schwarz. Estimating the dimension of a model. The annals of statistics, pages 461–464, 1978b.

J. P. Sexton, J. Montiel, J. E. Shay, M. R. Stephens, and R. A. Slatyer. Evolution of ecological niche breadth. Annual Review of Ecology, Evolution, and Systematics, 48:183–206, 2017.

S. Shalev-Shwartz and S. Ben-David. Understanding machine learning: From theory to algorithms. Cambridge university press, 2014.

J. K. Smith. Data transformations in analysis of variance. Journal of Verbal Learning and Verbal Behavior, 15(3):339–346, 1976.

J. Suman, A. Rakshit, S. D. Ogireddy, S. Singh, C. Gupta, and J. Chandrakala. Microbiome as a key player in sustainable agriculture and human health. Frontiers in Soil Science, 2:821589, 2022.

A. Susin, Y. Wang, K.-A. Lê Cao, and M. L. Calle. Variable selection in microbiome compositional data analysis. NAR Genomics and Bioinformatics, 2(2):lqaa029, 2020.

P. Vangay, B. M. Hillmann, and D. Knights. Microbiome learning repo (ml repo): A public repository of microbiome regression and classification tasks. Gigascience, 8(5):giz042, 2019.

A. Verster, N. Petronella, J. Green, F. Matias, and S. P. Brooks. A Bayesian method for identifying associations between response variables and bacterial community composition. PLoS Computational Biology, 18(7):e1010108, 2022.

L. Wang, Y. Xin, J. Zhou, Z. Tian, C. Liu, X. Yu, X. Meng, W. Jiang, S. Zhao, and Q. Dong. Gastric mucosa-associated microbial signatures of early gastric cancer. Frontiers in Microbiology, 11:1548, 2020.

C. Wen, Z. Zheng, T. Shao, L. Liu, Z. Xie, E. Le Chatelier, Z. He, W. Zhong, Y. Fan, L. Zhang, et al. Quantitative metagenomics reveals unique gut microbiome biomarkers in ankylosing spondylitis. Genome Biology, 18:1–13, 2017.

R. C. Wilhelm, H. M. van Es, and D. H. Buckley. Predicting measures of soil health using the microbiome and supervised machine learning. Soil Biology and Biochemistry, 164:108472, 2022.

S. Woodcock, T. P. Curtis, I. M. Head, M. Lunn, and W. T. Sloan. Taxa–area relationships for microbes: the unsampled and the unseen. Ecology Letters, 9(7):805–812, 2006.

T. Yatsunenko, F. E. Rey, M. J. Manary, I. Trehan, M. G. Dominguez-Bello, M. Contreras, M. Magris, G. Hidalgo, R. N. Baldassano, A. P. Anokhin, et al. Human gut microbiome viewed across age and geography. Nature, 486(7402):222–227, 2012.

W. Zhang, A. Liu, Z. Zhang, G. Chen, and Q. Li. An adaptive direction-assisted test for microbiome compositional data. Bioinformatics, 38(14):3493–3500, 2022.

A. Zheng and A. Casari. Feature engineering for machine learning: principles and techniques for data scientists. O’Reilly Media, Inc., 2018.

