## Supplemental Figures for "CODARFE: Unlocking the prediction of continuous environmental variables based on microbiome"

SUPPLEMENTARY MATERIAL

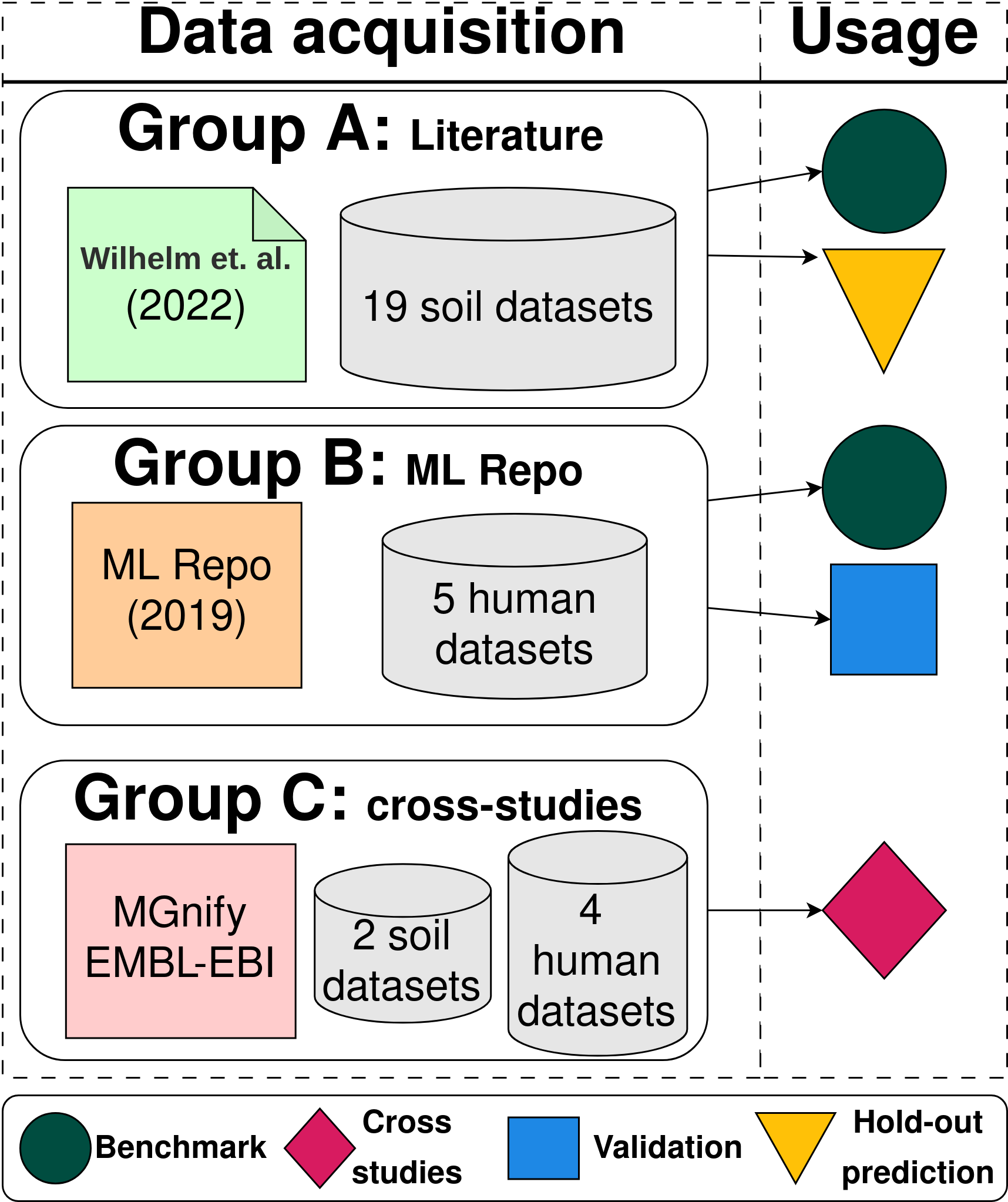

**Fig. S1. This flowchart shows the CODARFE’s data acquisition and usage. Group A: Literature comprises 19 soil health metrics extracted from the article [Wilhelm et al., 2022]. Group B: Machine Learning repository [Vangay et al., 2019], composed of five human disease indexes and linked to articles explaining taxa association. Group C: The MGnify database [Richardson et al., 2023] is formed by two sets: one of two soil datasets and four human health measures. Each group was used for one or more analyses, as depicted by the image.**

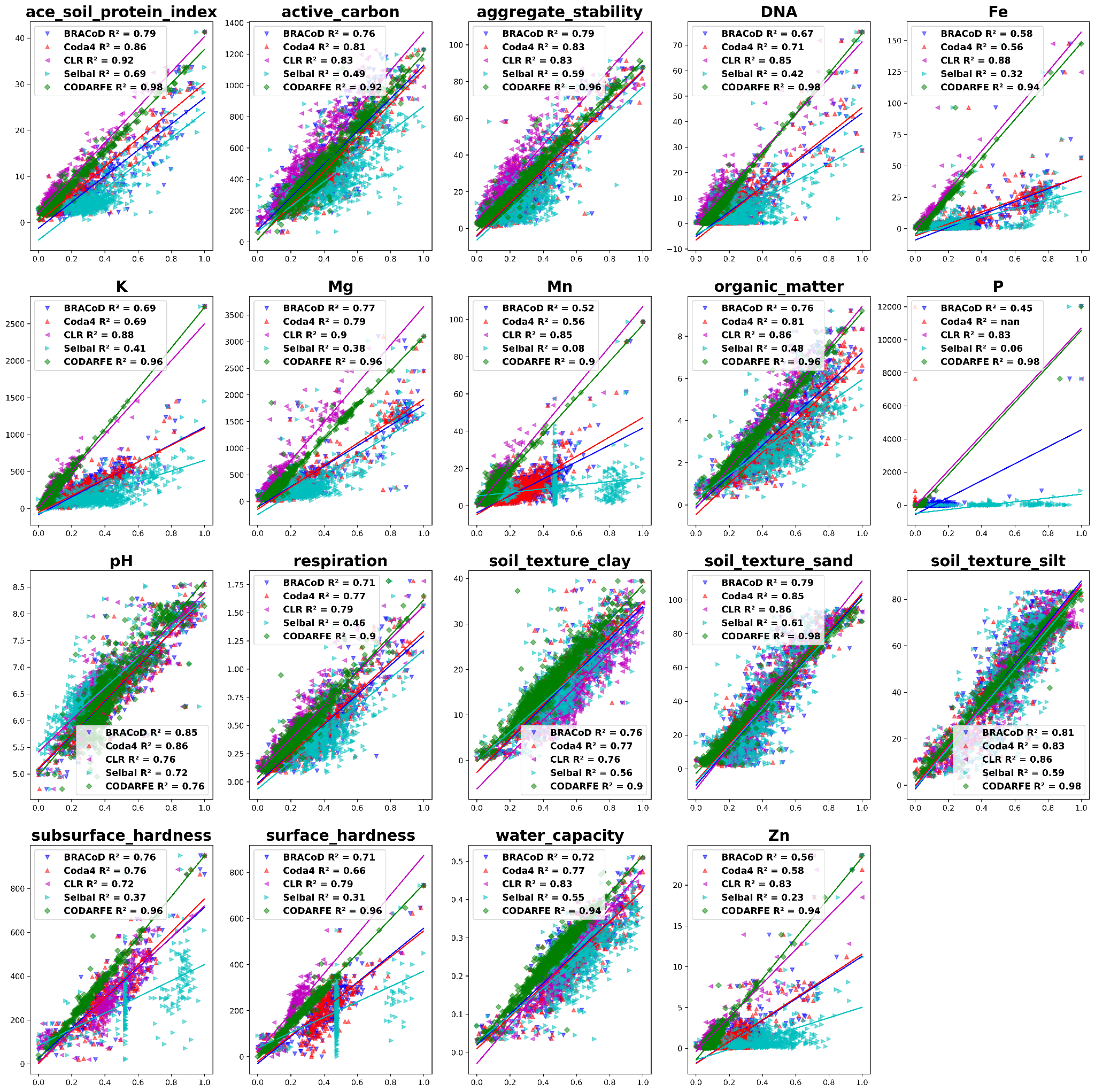

**Fig. S2. Complete distribution of all 19 soil health metrics across all tools. The y-axis represents the real value, while the x-axis represents the normalized (values between 0-1) prediction of each tool.**

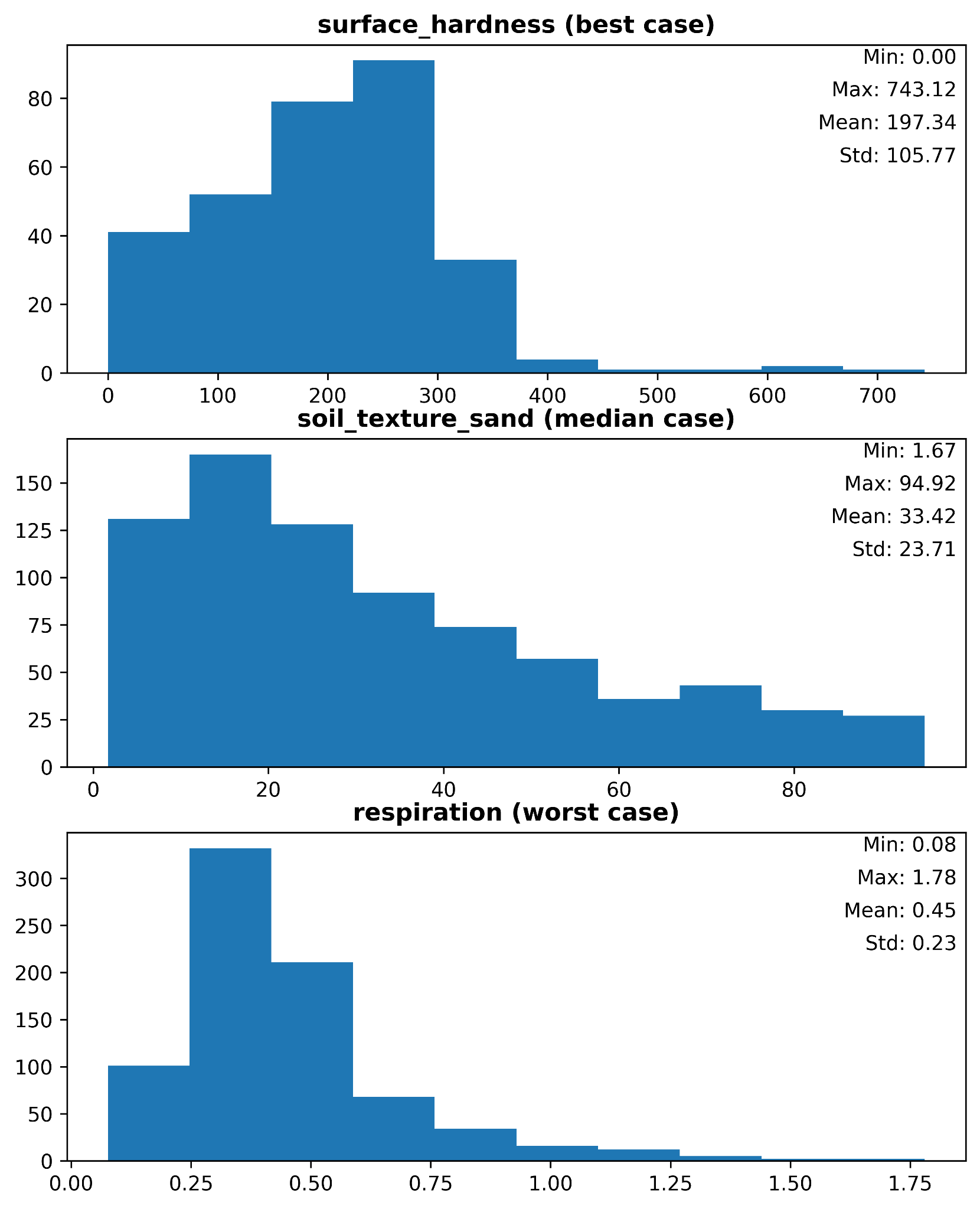

**Fig. S3. Distribution of the best, median and worst case of the hold-out validation process. The best case (surface_hardness) presents the highest amplitude, while the worst case (respiration) presents the lowest amplitude.**
